# Symptom observation underestimates co-infections: insight from viral and bacterial diseases in rice fields in Burkina Faso

**DOI:** 10.1101/2025.03.11.638810

**Authors:** Estelle Billard, Martine Bangratz, Abalo Itolou Kassankogno, Phonsiri Saengram, Abdoul Kader Guigma, Olivier Cotto, Gaël Thébaud, Issa Wonni, Drissa Sérémé, Nils Poulicard, Sebastien Cunnac, Mathilde Hutin, Eugénie Hébrard, Charlotte Tollenaere

## Abstract

Co-occurrence of multiple diseases and co-infection of individual plants by various pathogens have potential epidemiological and evolutionary implications. Based on previous information on the co-occurrence of the rice yellow mottle disease (caused by the rice yellow mottle virus, RYMV) and bacterial leaf streak (BLS, due to *Xanthomonas oryzae pv. oryzicola, Xoc*) in Burkina Faso, and experimental evidence of interactions between the pathogens causing these two diseases, we aimed to monitor the two pathogens more intensively in farmer’s rice fields. To this purpose, we selected fields showing both types of symptoms to maximize the chance of observing co-infections at the plant scale. We performed observations and sampling in two sites over two consecutive years. Over a global dataset of 1666 samples, 1341 were symptomatic. Although the sampling design aimed to observe co-infections, only 37 of these samples (2.8%) were annotated as presenting both yellow mottle and BLS symptoms. The samples were then subjected to a newly designed molecular detection test that specifically amplifies both the virus (RYMV) and the bacteria (*Xo*). This revealed that 166 samples, i.e. 12.4% of symptomatic samples, were co-infected by RYMV and *Xo*, hence showing that symptom observation in the field greatly underestimates co-infection levels. Combining these data with a previously published dataset, we estimated that up to 1-4% of all plants in disease hotspots are simultaneously infected by the two pathogens. Further research on multiple infections would benefit from longitudinal surveys over the crop growing season rather than such a cross-sectional study.

## MANUSCRIPT

The co-circulation of multiple diseases at the local scale may result in individual plants being infected by different pathogens, i.e. co-infections. In this context, pathogens may interact within plants, potentially affecting the epidemiological and evolutionary trajectories of pathogen populations (Tollenaere et al. 2016). This phenomenon has recently gained attention, particularly for plant viruses (Alcaide et al. 2020), some of which cause well-known disease complexes, e.g. maize lethal necrosis (Wamaitha et al. 2018), or are involved in striking phenotypes resulting from synergistic or antagonistic interactions (Moreno and López-Moya 2019; Syller 2012). However, epidemiological studies following up different diseases in space and time still remain scarce (Jokinen et al. 2023; Maclot et al. 2023; Pagan et al. 2010), especially when considering pathogens from different kingdoms.

Major rice diseases in West Africa include the rice yellow mottle disease, caused by the rice yellow mottle virus, RYMV (Hébrard et al. 2021), and bacterial leaf streak, BLS, caused by *Xanthomonas oryzae* pv. *oryzicola, Xoc* (Wonni et al. 2014). These two diseases have been shown to co-circulate in the same rice fields in western Burkina Faso (Barro et al. 2021; Tollenaere et al. 2017), and experimental work showed that the RYMV-*Xoc* co-infection can lead to asymmetric interactions, with lower viral loads but stronger bacterial symptoms, compared to the mono-infection conditions (Tollenaere et al. 2017).

In this study, we aim to further characterize the co-circulation of these two diseases in hotspot areas and to refine the estimate of RYMV-*Xoc* co-infection levels in rice fields. To this end, we designed a sampling scheme that focused on sites and fields where the two diseases co-occurred. Two study sites in western Burkina Faso were selected based on existing data on the epidemiology of yellow mottle disease and bacterial leaf streak (see Barro et al. 2021; doi.org/10.23708/8FDWIE): the rainfed lowland of Badala, and the irrigated perimeter of Banzon (Fig. S1). Studied fields (small rice plots; ca. 0.05 ha) were visited over two consecutive years (2021 and 2022), during the main cropping season and at the same stage of rice growth (see: doi.org/10.23708/QB8BOO). Within each site, a preliminary survey identified the fields to be studied, preferably selecting fields where symptoms of both diseases could be observed. Twenty fields were visited in 2021, and 25 in 2022 (Fig. 1).

**Figure 1:**
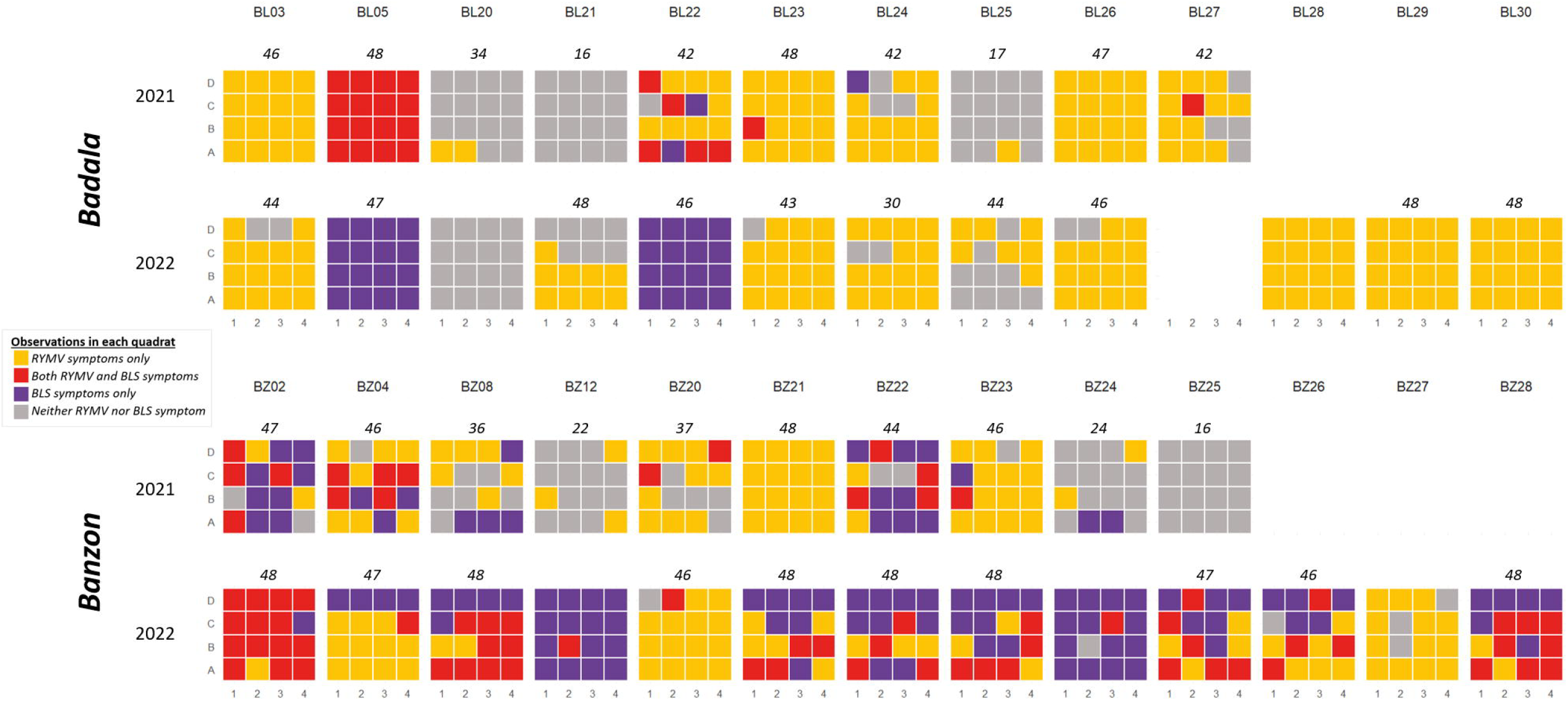
Observations of the symptoms of the two rice diseases (yellow mottle disease and bacterial leaf streak) in each of the 16 quadrats of the surveyed fields in western Burkina Faso. The internal squares represent quadrats in the visited fields of the two sites (Badala and Banzon) and the two studied years (2021 and 2022). The number of samples analyzed is also indicated for the 40 fields for which samples were analyzed using molecular detection tools.

In each field, we set up a grid of 16 quadrats (Fig. 1). Sampling was conducted using a stratified random method, with three samples collected per quadrat. For each individual rice plant sampled, three leaves were collected in a paper envelope. Samples from the same plot were then grouped in a labeled plastic bag containing silica gel to allow the collected leaves to dry quickly. The information associated with each sample was collected using the KOBO collect application (www.kobotoolbox.org). For each field, we asked the farmers for permission to visit their fields and collect samples. All samples collected are the property of INERA.

A subset of the samples (10 fields per site per year; only one sample in the quadrats where no symptoms were observed) were subjected to nucleic acid extractions using the Plant RNA-DNA Purification Kit (Norgen Biotek, Thorold, Canada) in a 96-well format. We immersed leaf samples (∼35-50 mg) in liquid nitrogen and disrupted them in 2-mL tubes containing steel beads using Tissue Lyser (Qiagen, Hilden, Germany) for 30 s at 30 Hz. We then followed the manufacturer’s recommendations for extraction (see doi.org/10.23708/QB8BOO). We used a newly designed method to simultaneously detect RYMV (based on CP gene amplification, see Pinel et al. 2000) and *Xanthomonas oryzae* (using the primers designed by Lang et al. 2010). The reliability of the detection method and the set-up of the protocol were assessed beforehand by comparing simplex and multiplex PCR, and evaluating the best annealing temperature to optimize PCR amplification. Reverse transcription (RT) was performed on 7.5µL of nucleic acid extract using 0.1µM of primer RYMV_M and 200U of M-MLV reverse transcriptase (Thermo Fisher, Waltham, USA). A mixture of nucleic acid extract (4µL) and RT product (1µL) was subjected to PCR amplification using Firepol Master Mix (Solis Biodyne, Tartu, Estonia), and the following primers: RYMV_M, RYMV_III, Xo_3756F and Xo_3756R. The initial denaturation (95°C, 15 min) was followed by 30 cycles (95°C for 30s, 58°C for 15s, and 72°C for 45s) and a final extension at 72°C for 5 min. Specific amplicons of 331nt (*Xo*) and 765nt (RYMV) were visualized on a 2% agarose gel.

We obtained a dataset of 1666 samples, of which 893 were reported as symptomatic for RYMV only, 411 for BLS only, 37 for both diseases and 325 without symptoms (Table 1; Fig. 2; doi.org/10.23708/QB8BOO). Molecular detection revealed the presence of only the virus in 761 samples, only the bacteria *Xo* in 384 samples, both RYMV and *Xo* in 176 samples and no pathogen in the remaining 345 samples (Table 1).

**Table 1:** Number of rice samples for the different categories of observed symptoms in the field and molecular detection outcome for the two sites (Badala and Banzon) and years (2021 and 2022), as well as overall (i.e. for all sites and years). The observed symptoms correspond to four categories: yellow mottle symptoms specific to the rice yellow mottle virus (RYMV only), both yellow mottle and bacterial leaf streak symptoms observed on the same leaves (RYMV+BLS), bacterial leaf streak (BLS only) symptoms specific of *Xanthomonas oryzae* (*Xo*) pv. *oryzicola*, and no symptom observed on this plant in the field. Molecular detection also results in four categories: RYMV detection only, both RYMV and *Xo* detected on the same sample, *Xo* detection only, and neither pathogen detected. The percentages are relative to the number of plants in each symptom category.

| Site | Year | Symptoms | Molecular detection result |  |  |  |  |
| --- | --- | --- | --- | --- | --- | --- | --- |
|  |  |  | RYMV only | RYMV+*Xo* | *Xo* only | None |  |
| Badala | 2021 | RYMV only | 209 (86.4%) | 23 (9.7%) | 0 (0%) | 10 (3.9%) | **242** |
|  |  | RYMV+BLS | 1 (25.0%) | 3 (75.0%) | 0 (0%) | 0 (0%) | **4** |
|  |  | BLS only | 2 (6.3%) | 6 (18.8%) | 21 (65.6%) | 3 (9.4%) | **32** |
|  |  | None | 3 (2.9%) | 1 (1.0%) | 8 (7.7%) | 92 (88.5%) | **104** |
|  |  |  | **215** | **33** | **29** | **105** | **382** |
|  | 2022 | RYMV only | 233 (75.6%) | 7 (2.3%) | 1 (0.3%) | 67 (21.8%) | **308** |
|  |  | RYMV+BLS | 0 (0%) | 0 (0%) | 0 (0%) | 0 (0%) | **0** |
|  |  | BLS only | 0 (0%) | 6 (6.3%) | 75 (78.9%) | 14 (14.7%) | **95** |
|  |  | None | 6 (14.6%) | 0 (0%) | 0 (0%) | 35 (85.4%) | **41** |
|  |  |  | **239** | **13** | **76** | **116** | **444** |
| Banzon | 2021 | RYMV only | 120 (84.5%) | 17 (12.0%) | 1 (0.7%) | 4 (2.8%) | **142** |
|  |  | RYMV+BLS | 0 (0%) | 0 (0%) | 0 (0%) | 0 (0%) | **0** |
|  |  | BLS only | 3 (3.7%) | 27 (33.3%) | 46 (56.8%) | 5 (6.2%) | **81** |
|  |  | None | 20 (14.0%) | 8 (5.6%) | 42 (29.4%) | 73 (51.0%) | **143** |
|  |  |  | **143** | **52** | **89** | **82** | **366** |
|  | 2022 | RYMV only | 142 (70.6%) | 33 (16.4%) | 19 (9.5%) | 7 (3.5%) | **201** |
|  |  | RYMV+BLS | 15 (45.5%) | 7 (21.2%) | 7 (21.2%) | 4 (12.1%) | **33** |
|  |  | BLS only | 6 (3.0%) | 37 (18.2%) | 149 (73.4%) | 11 (5.4%) | **203** |
|  |  | None | 1 (2.7%) | 1 (2.7%) | 15 (40.5%) | 20 (54.1%) | **37** |
|  |  |  | **164** | **78** | **190** | **42** | **474** |
| Overall | 2021-2022 | RYMV only | 704 (78.8%) | 80 (9.0%) | 21 (2.4%) | 88 (9.9%) | **893** |
|  |  | RYMV+BLS | 16 (43.2%) | 10 (27.0%) | 7 (18.9%) | 4 (10.8%) | **37** |
|  |  | BLS only | 11 (2.7%) | 76 (18.5%) | 291 (70.8%) | 33 (8.0%) | **411** |
|  |  | None | 30 (9.2%) | 10 (3.1%) | 65 (20.0%) | 220 (67.7%) | **325** |
|  |  |  | **761** | **176** | **384** | **345** | **1666** |

**Figure 2:**
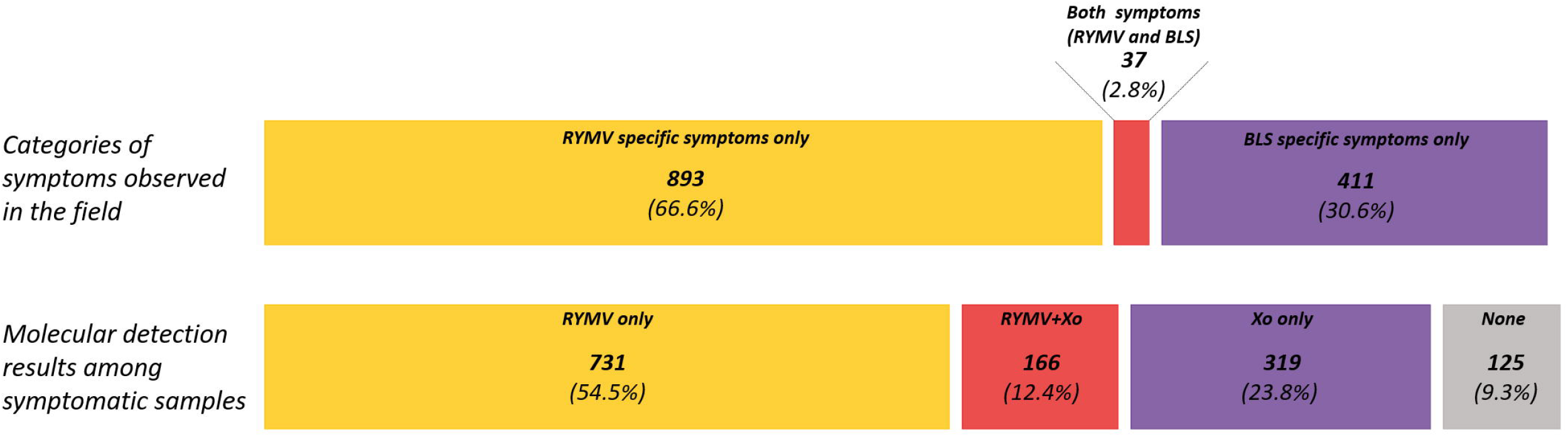
Distribution of the symptom categories and molecular detection results among the 1341 symptomatic samples analyzed in this study. In yellow are represented the samples infected by the RYMV (based on symptoms above and specific molecular detection below). In purple are the samples presenting BLS symptoms (top) and *Xanthomonas oryzae* (*Xo*) positive in molecular detection (bottom). In red are the samples with both types of symptoms (top) and positive for both pathogens according to molecular results (bottom). In grey are the samples where no pathogen could be detected (bottom). This analysis is performed on symptomatic samples only (1341 samples out of 1666 in total); the non-symptomatic samples (325 samples) are not represented.

Comparing the molecular detection results with symptom observations, we noticed that among the 893 samples annotated as symptomatic for RYMV only, 784 (87.8%) were RYMV-positive in molecular detection (including the RYMV-*Xo* samples, Fig. 3a). Similarly, of the 411 samples annotated as symptomatic for BLS only, 367 (89.3%) were *Xo*-positive (also including the RYMV-*Xo* samples, Fig. 3a). Therefore, we considered the symptom observation carried out in the field as globally accurate. However, when the dataset is divided according to the site and year (Table 1 and Fig. 3b), these frequencies are more variable for RYMV (ranging from 77.9% to 96.5%), than for BLS (84.4-91.6%). It should be noted that the molecular method could not distinguish between *Xoc* and *Xoo* (pv. *oryzae*, which also infects rice but causes different symptoms). However, this congruence with the symptom observations, as well as further analysis of these samples (unpublished), suggests the local absence (or very low prevalence) of *Xoo*. On the other hand, for RYMV, the relationship between symptoms and molecular diagnostic is less clear for Badala in 2022, which may be due to the circulation of another virus (unpublished) and subsequent partial local confusion of symptoms.

**Figure 3:**
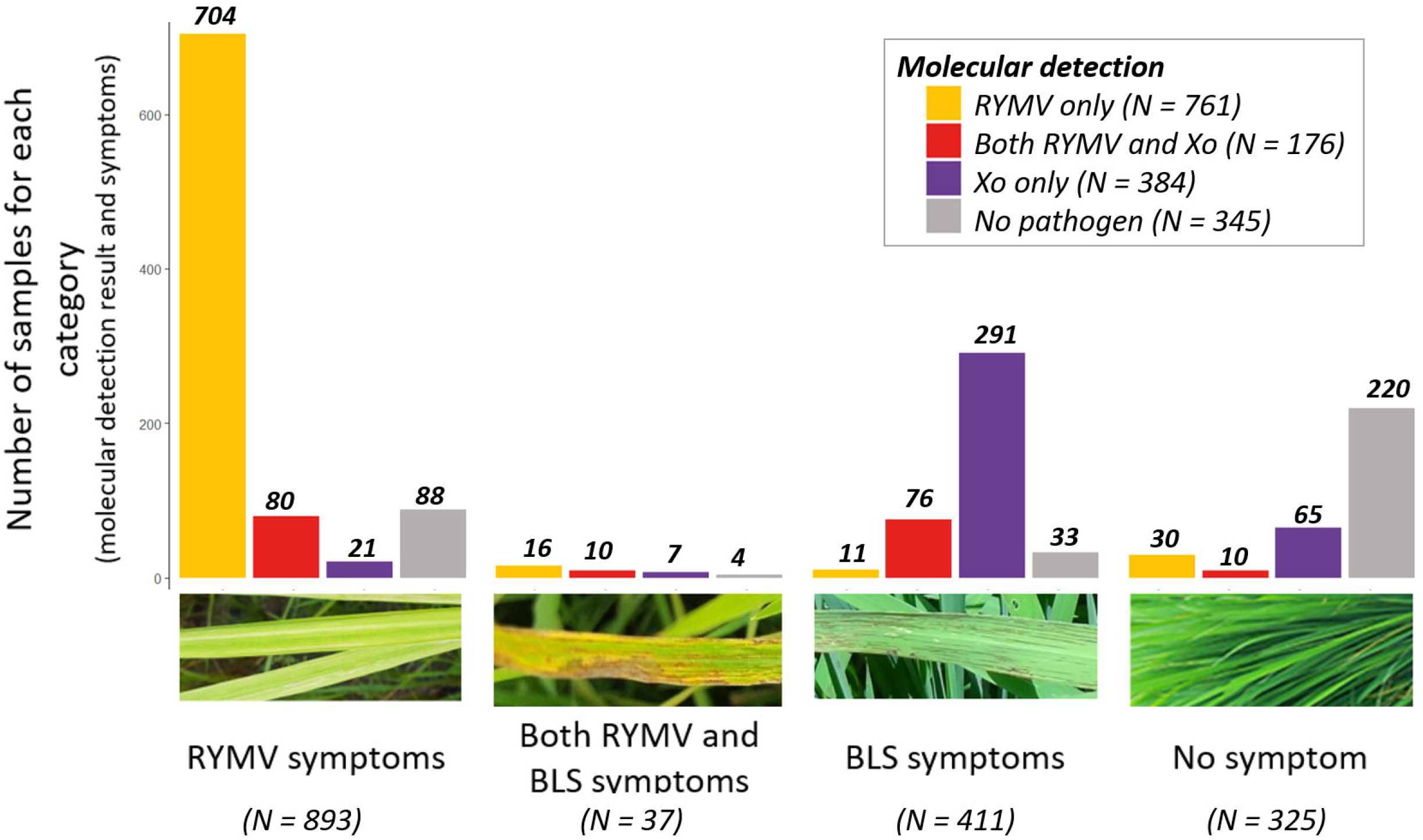
Distribution of the 1666 analyzed rice samples into the different categories of observed symptoms (yellow mottle, BLS, both on the same plant, no symptom), and the result of the specific detection of RYMV and *Xo* rice pathogens. a, repartition of the samples over the whole dataset (two sites and two years); b, subdivision of the dataset for each site (Badala on the left and Banzon on the right) and year (2021 and 2022). The images illustrate the symptom categories assigned in the field.

Considering multiple infections based on field observation, only 37 samples (2.8% of symptomatic samples) presented the two types of symptoms (Fig. 2 and 3a). Among all collected symptomatic samples (1341 samples, Table 1 and Fig. 2), 166 (12.4%) tested positive for both RYMV and *Xo* infections. The same patterns were found at the site and year level (Fig. 3b). We note that among the 893 samples annotated as “RYMV symptoms only”, 80 were found positive to both RYMV and *Xo* (9.0%, Fig. 3a, ranging from 2.3% in Badala 2022 to 16.4% in Banzon 2022, Fig. 3b). Among the “BLS-only” symptomatic samples (411), both pathogens were detected in 76 samples (18.5%, ranging from 6.3% in Badala 2022 to 33.3% in Banzon 2021).

While our sampling was focused on the fields presenting both diseases, we used data collected in 2016-2019, from six sites in western Burkina Faso (Barro et al. 2021) to infer global co-infection levels. In this previous study where the sampling design is not biased towards symptoms, the global (6 sites and 4 years) frequency of fields with the two diseases (based on symptoms) was 26 out of 179 fields (14.5%) studied over the 2016-2019 period. Locally, this frequency was of 22.2% (6/27) in Badala, while it reached 42.4% (14/33) in Banzon. Within these fields where the two diseases co-circulate, we obtained a frequency of symptomatic samples (specific for at least one of the two diseases: RYMV or BLS) of 29.2% (28/96 samples) in Badala and 45.5% (102/224) in Banzon. In the present study, focusing on the fields where symptoms of both diseases were observed (see Fig. 1: 5 fields in Badala in 2021, 7 fields in Banzon in 2021 and 10 fields in Banzon in 2022), and on symptomatic samples within these fields, we counted 28 samples as positive for the two pathogens out of 183 (15.3%) in Badala and 121/609 (19.9%) in Banzon. Globally, the frequency of RYMV-*Xo* co-infection over all rice plants can therefore be estimated to be 1.0% in Badala and 3.8% in Banzon.

Unfortunately, our sampling design did not allow to test for interactions in the field, e.g. using the approach of Hilker et al. (2017). However, detecting pathogen-pathogen interactions from field data is not straightforward (Hamelin et al. 2019), and discrepancies between experimental and field data have been reported (see e.g. Ezenwa and Jolles 2015; Halle et al. 2024). Interactions may be affected by environmental factors, and even depend on the genotypes of the protagonists (Seppala et al. 2009). As multiple genotypes of both RYMV (Billard et al. 2023) and *Xoc* (Wonni et al. 2014) co-exist in the studied sites, the perspectives include characterizing the genetic diversity of the two pathogens for this specific sampling. Furthermore, cross-sectional data (capturing the situation at one timepoint) may not be sufficient to analyze interactions, as the order of infections (priority effect) may shape the patterns (Powell-Romero et al. 2024). Longitudinal surveys would be necessary to unravel the co-circulation of multiple pathogens, a current challenge for epidemiology (Jokinen et al. 2023) and crop protection (Hilker et al. 2024).

## ACKNOWLEDGEMENTS

This work was carried out thanks to the facilities of the “International joint Laboratory LMI PathoBios: Observatory of plant pathogens in West Africa: biodiversity and biosafety” (www.pathobios.com). We are grateful to Zougrana Sylvain, Kafando Aida, Sanou Leïla Pulchérie, Nana Abdoulaye, Ouattara Bakary and Zongo Bowendsom who contributed to the sampling in southwestern Burkina Faso, and to the rice farmers from the two study sites (Badala and Banzon) for their collaboration. We also thank Félicité Kaboré for her contribution to implement KOBO collect application. This work has been publicly funded by the ANR (the French National Research Agency) under the project EVCOPAR (ANR-20-CE35-0004-01).

## SUPPLEMENTARY INFORMATION

**Supplementary Figure 1:**
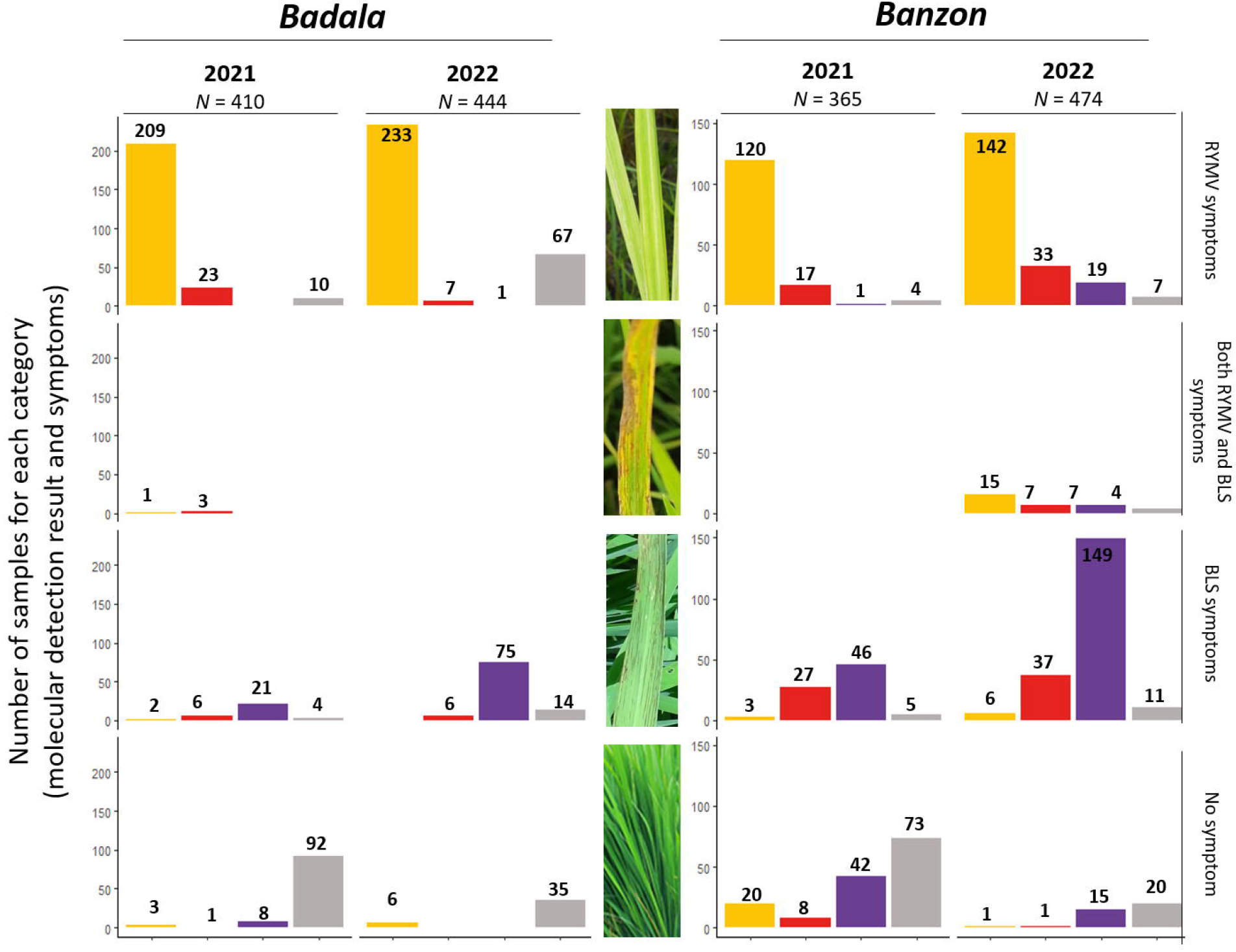
Map showing the location of the studied fields in western Burkina Faso.

