## Supplementary figures and images for "Symptom observation underestimates co-infections: insight from viral and bacterial diseases in rice fields in Burkina Faso"

### Supplementary Figure 1

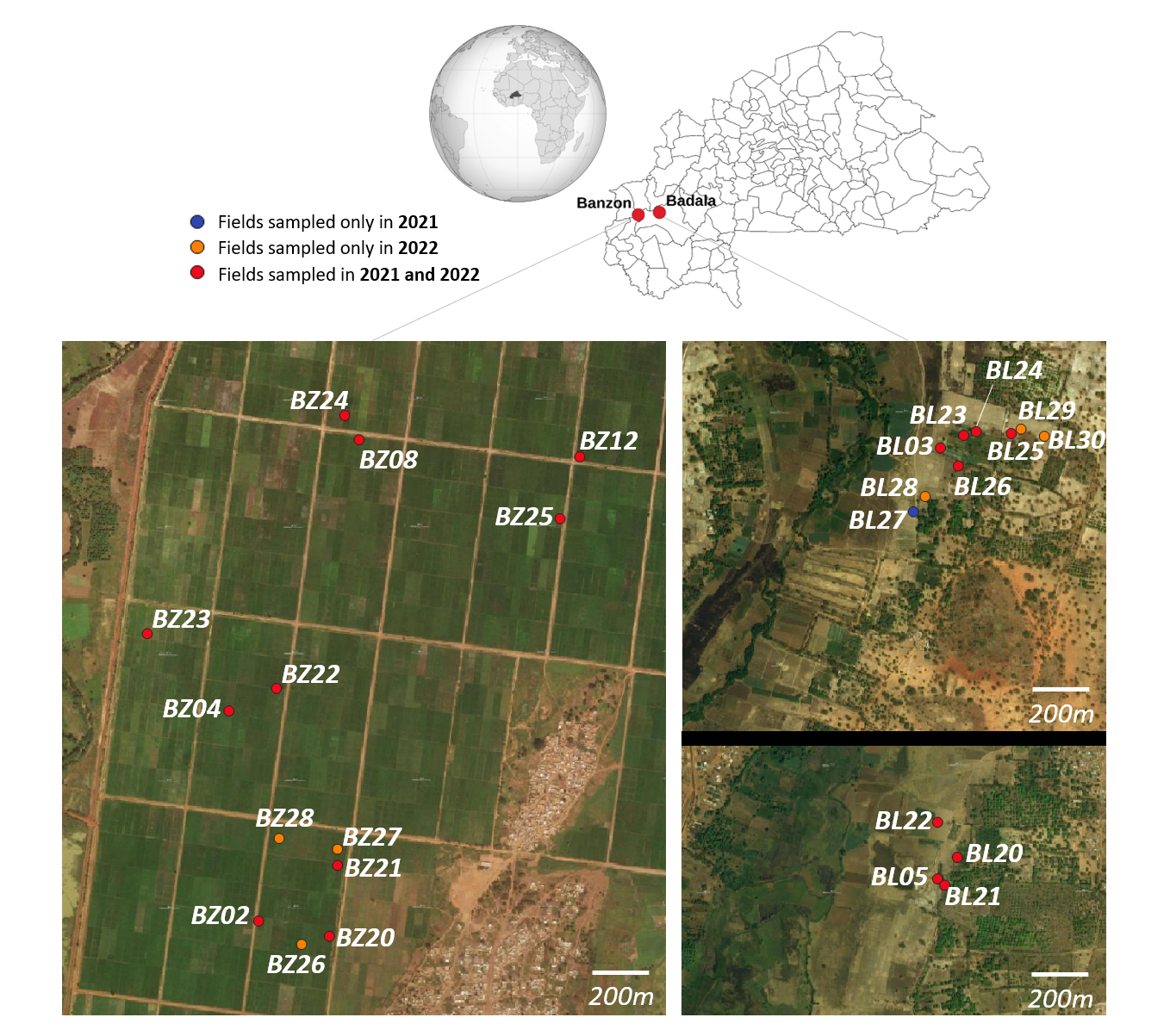
